# Exploring the interaction of inhibitor ‘YM-1’ with *Plasmodium falciparum* HSP70s through *in silico* methods

**DOI:** 10.1101/2023.08.21.554099

**Authors:** Bikramjit Singh, Satinder Kaur, Chanchal Nainani, Komalpreet Kaur Sandhu, Vipul Upadhyay, Rachna Hora, Prakash Chandra Mishra

**Author notes:** Correspondance: Prakash Chandra Mishra, Extn: 3220. Equal contribution (Bikramjit Singh, Satinder Kaur).

## Abstract

YM-1 belongs to the rhodacyanine class of the 70kDa heat shock protein (Hsp70) inhibitors. It antagonizes nucleotide exchange factor (NEF) ‘BAG3’ to block the critical ATPase cycle of HSP70 chaperones thereby disabling them. The anti-cancer potential of this small molecule derivative of MKT-077 is attributed to its improved solubility and stability over its parent. In the current study, we have used bioinformatics tools including molecular docking to study the binding strength of YM-1 for *Plasmodium falciparum* (Pf) HSP70s. Of the four PfHSP70 homologs, the mitochondrial counterpart was predicted to bind most efficiently with YM-1 at a previously reported inhibitor binding site. The interaction of YM-1 with mitochondrial PfHSP70 was significantly better than with any of the other Pf homologs. Further analysis of this PfHSP70-YM-1 complex helped us to predict not just the types of interactions, but also specific residues of the protein involved in inhibitor engagement. Information reported in this article is important from the standpoint of design and development of newer rhodacyanine based drugs against malaria.

## Introduction

Heat shock proteins of 70 kDa (Hsp70) are elevated in various cancers with their levels being correlated with poor prognosis in patients [1]. These function to control cellular senescence, apoptosis and host immune responses in cancerous cells possibly through their ability to handle stress by protein refolding, prevention of aggregation, protein degradation *etc*. Hsp70s are chaperones that interact with a variety of ancillary proteins like co-chaperones Hsp40, nucleotide exchange factors (NEF) and Hsp90s to facilitate protein homeostasis. While Hsp40s are positive regulators of the ATPase action of Hsp70s, NEFs modulate ATP binding of Hsp70 at its N-terminal nucleotide binding domain (NBD). Several classes of small molecule inhibitors have been designed to work against the Hsp70s in an attempt to develop anti-cancer drugs. Such drugs are also expected to be active in neurodegenerative and infectious diseases that induce stress [2], [3].

Rhodacyanines form a category of inhibitors that block the functional ATPase cycle of HSP70s through their binding within the NBD at allosteric sites [4]. This results in the molecule being locked in its ADP bound state that disallows substrate release and initiation of a fresh ATPase cycle by Hsp70. MKT-077, a member of the rhodacyanine class of molecules failed to pass Phase I clinical trials owing to its short half-life and renal toxicity. Soon its analogues *viz*. YM-1, JG-83, JG-84, JG-98, JG-231 and JG-294 *etc*. [4], [5] gained impetus as they had improved stability post administration and were more found more effective at targeting a variety of cancerous cells. YM-1 is a stable and soluble MKT-077 analogue with heightened and specific cytotoxic action against cancerous cells [6]. While parent MKT-077 accumulates in the mitochondria, YM-1 localizes to the cell cytosol. Brief treatment of drug resistant breast cancer cells with YM-1 was found to re-sensitize them to tamoxifen.

YM-1 blocks the interaction between HspA1A (an inducible human Hsp70) and BAG3 (Bcl-2-associated AnthanoGene-3), one of the many NEFs in humans [7]. This in turns inhibits Hsp70 driven Src signalling. BAG proteins contain a 100 amino acid long BAG domain that effects their nucleotide exchange function [8]. These also function as adaptor molecules that form complexes with various signalling proteins or Hsp70s via multiple motifs to obtain diverse functionalities. Of the six human BAG proteins, BAG3 is the largest that is also reported to bind the substrate binding domain (SBD) of HSPA8 (constitutively expressed cytosolic human Hsp70) through a non-BAG domain. BAG3 is a critical player in destruction of polyubiquitinated client proteins by autophagy [9].

Hsp70s are also modulated by binding to the middle domain of Hsp70 interacting proteins (HIP), which behave as anti-exchange factors. HIPs are therefore antagonistic in action to BAGs, stabilize the ADP bound state of Hsp70s and cause protein folding in the absence of NEF [10]. YM-1 is hence believed to act similar to HIP so as to increase the apparent affinity of Hsp70 for its substrates by confining it in its ADP bound open state. Once bound to client proteins for a long period, the C-terminus of HIP (CHIP), an E3 ubiquitin ligase mediates their ubiquitination and degradation.

It is evident that the Hsp70 family of proteins are structurally conserved across different organisms and bind a variety of modulators to bring about different cellular functions. Rhodacyanine inhibitors rhodamine and MKT-077 have earlier shown anti-malarial potential against cultured *Plasmodium falciparum* (Pf) parasites [3]. Pf expresses a total of four HSP70s: PfHSP70-1 that localizes to the parasite nucleus and cytosol, PfHSP70-2 that homes in endoplasmic reticulum, PfHSP70-3 is present in mitochondria and PfHSP70-x gets exported to parasite infected erythrocytes. Since the MKT-077 derivative ‘YM-1’ binds to multiple Hsp70s in order to inhibit their interaction with BAG3, we performed *in-silico* analysis to identify the YM-1 binding pocket on all four PfHSP70s.

## Materials and methods

### Sequences and structures

The three-dimensional structures of NBDs of PfHSP70-2 and PfHSP70-x were downloaded from the protein data bank (PDB) database (5UMBand 6S02 respectively) **[11], [12]**. Homology modeling for the structures of NBDs of PfHSP70-1 and PfHSP70-3 had earlier been done in our laboratory by SwissModel and validated by Ramachandran plot analysis and ERRAT **[13]**. Structure of YM-1 inhibitor was obtained from PubChem database **[14]**.

### Identifying the probable binding region of YM-1 on PfHSP70s

We have analysed the interaction of YM-1 at the ADP and HEW binding sites on PfHSP70s. The ADP binding site was determined from the three-dimensional structure of ADP bound PfHSP70-2 NBD (5UMB). The HEW (2-amino 4 bromopyridine) binding site was determined on the basis of the crystallographic structure of inhibitor bound PfHSP70-x (7OOG). LigPlot analysis was performed on these structures downloaded from PDB to map the binding regions for ADP and HEW **[15], [16]**. A list of interacting residues present in both the binding pockets were deduced for other PfHSP70 homologs by pairwise sequence alignment **[17]**.

### *In silico* docking and analysis of complexes

UCSF chimera was used to prepare the protein and ligand structures for the docking experiments **[18]**. Grid boxes were made on the prepared structures keeping in mind the predicted ligand binding sites on PfHSP70 homologs. After this, Autodock Vina plugin in Chimera was used to dock YM-1 on ADP and HEW binding sites of all four PfHSP70s to obtain eight sets of complexes **[19]**. For each set, the best one was selecting on the basis of binding energies. The best complex (least binding energy) was analyzed by using LigPlot to predict the interacting amino acids.

## Results and discussion

### Structures of proteins and YM-1 for docking

YM-1 is a derivative of MKT-077, which is reported to interact with human PfHSP70s **[20]**. YM-1 is considered superior to MKT-077 owing to its improved stability and solubility **[6]**. In the present study, we have predicted the YM-1 binding pockets on all four PfHSP70 homologs. For this, structures of NBDs of PfHSP70-x and PfHSP70-2 were downloaded from PDB (6S02 and 5UMB respectively) (Fig 1a-b). Structures of the corresponding domains for PfHSP70-1 and PfHSP70-3 were derived by using homology modeling in automatic mode of SwissModel **[21]** (Fig 1c-d). These modeled structures were validated by Ramachandran plot analysis, where none of the amino acids were found in disallowed regions (data not shown). The structure of YM-1 was downloaded from PubChem (CID 10895023) in SDF format (Fig 1e) **[14]**.

**Figure 1:**
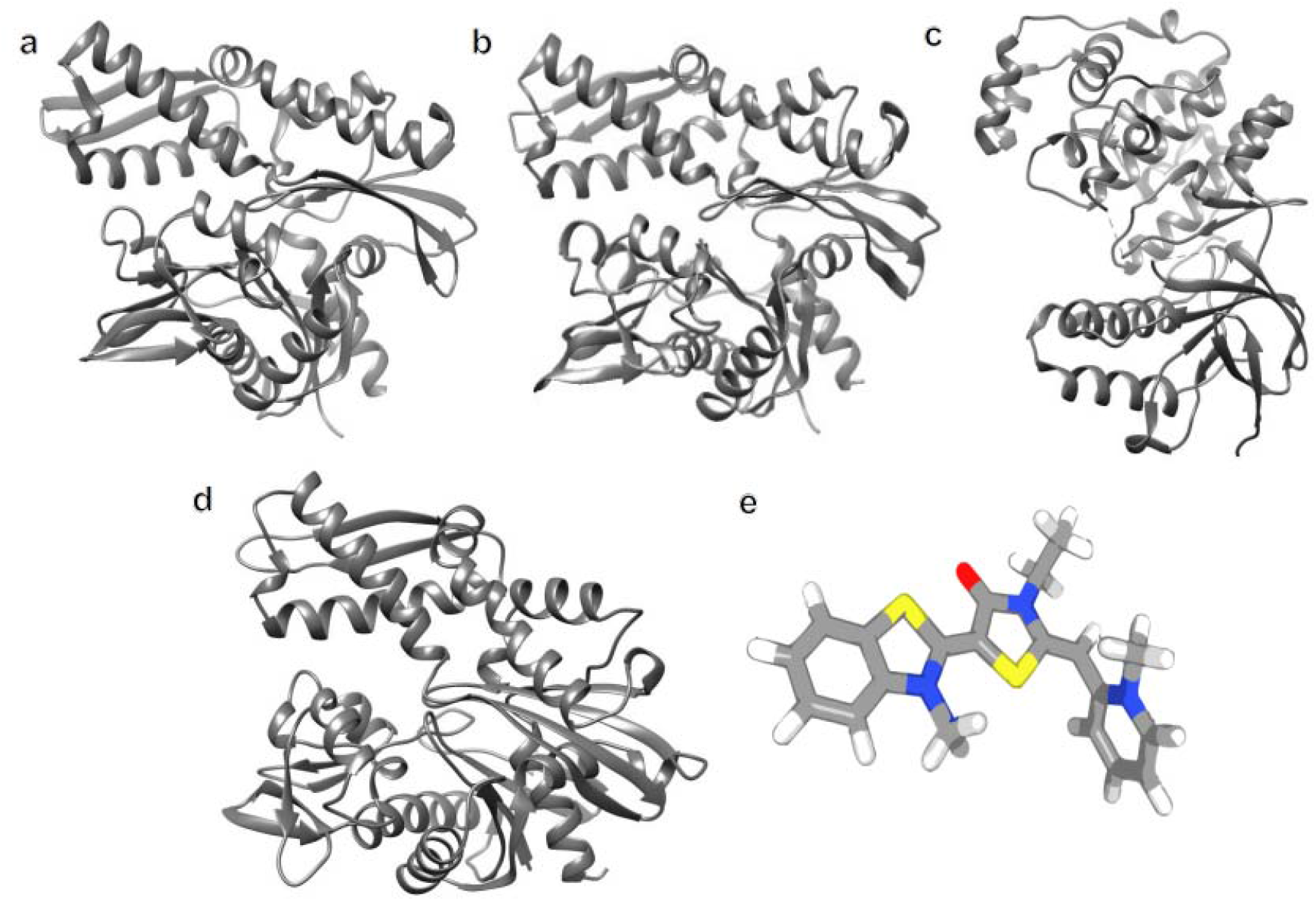
Protein and ligand structures. Ribbon diagrams of three-dimensional structures of nucleotide binding domains (NBD) of PfHSP70 homologs of a. PfHSP70-x (PDB ID: 6S02) b. PfHSP70-2 (PDB ID: 7OOG) c. PfHSP70-1 d. PfHSP70-3. c & d were obtained through molecular modelling by SwissModel c. PfHSP70-1 e. Structure of YM-1 (SDF format) obtained from PubChem (CID 10895023).

### *In silico* docking of YM-1 on PfHSP70s

UCSF Chimera with AutoDock Vina plugin was used to dock YM-1 on two different binding sites present on all PfHSP70 homolog structures **[18]**. The two sites were the ADP binding and the inhibitor ‘HEW’ binding pockets of PfHSP70s **[22]**. These sites were derived by performing LigPlot analysis of crystallographic three dimensional structures PfHSP70-2-ADP (PDB ID: 5UMB) and PfHSP70-x-HEW (7OOG) **[15]** (Table 1). ADP and HEW binding sites on the other homologs were predicted through pairwise sequence alignment (Table 1). This information was used to generate grid boxes on all the PfHSP70 homologs for outlining protein regions to be used for the docking experiments (Fig. 2). The centre and size of the gridlines are provided (Table. S1). The grid box dimensions for the HEW binding sites of PfHSP70-1 and PfHSP70-x were borrowed from Singh *et al* **[13]**. Molecular docking of each protein-ligand pair for either binding site was performed to obtain docked complexes.

**Table 1:** Amino acids of PfHsp70 homologs showing ADP or HEW binding. Residues of PfHSP70-2 and PfHSP70-x that interact with ADP or HEW respectively obtained from LigPlot analysis of crystallographic data are highlighted in bold. Residues of other homologs predicted to bind with ADP/HEW through pairwise alignment are also tabulated.

| Protein homologue | ADP binding residues | HEW binding residues |
| --- | --- | --- |
| **PfHSP70-1** | Asp21, Tyr27, Thr25, Gly214, Lys284, Glu279, Gly352, Arg355, | Arg83, His100, Gly215, His239, Asp244, Glu243 |
| **PfHSP70-2** | Thr38, Tyr39, Gly224, Lys293, Gly361, Ser297 | Arg96, Gly225, His249, Glu253, Asp254 |
| **PfHSP70-3** | Thr12, Tyr13, Gly198, Lys267, Ser271, Gly335, | Arg70, Gly199, His223, Glu227, Asp228 |
| **PfHSP70-x** | Asp39, Tyr43, Tyr44, Gly232, Gly299, Lys302, Gly370, Arg373 | Arg101, His118, Gly233, Glu261, Asp262 |

**Figure 2:**
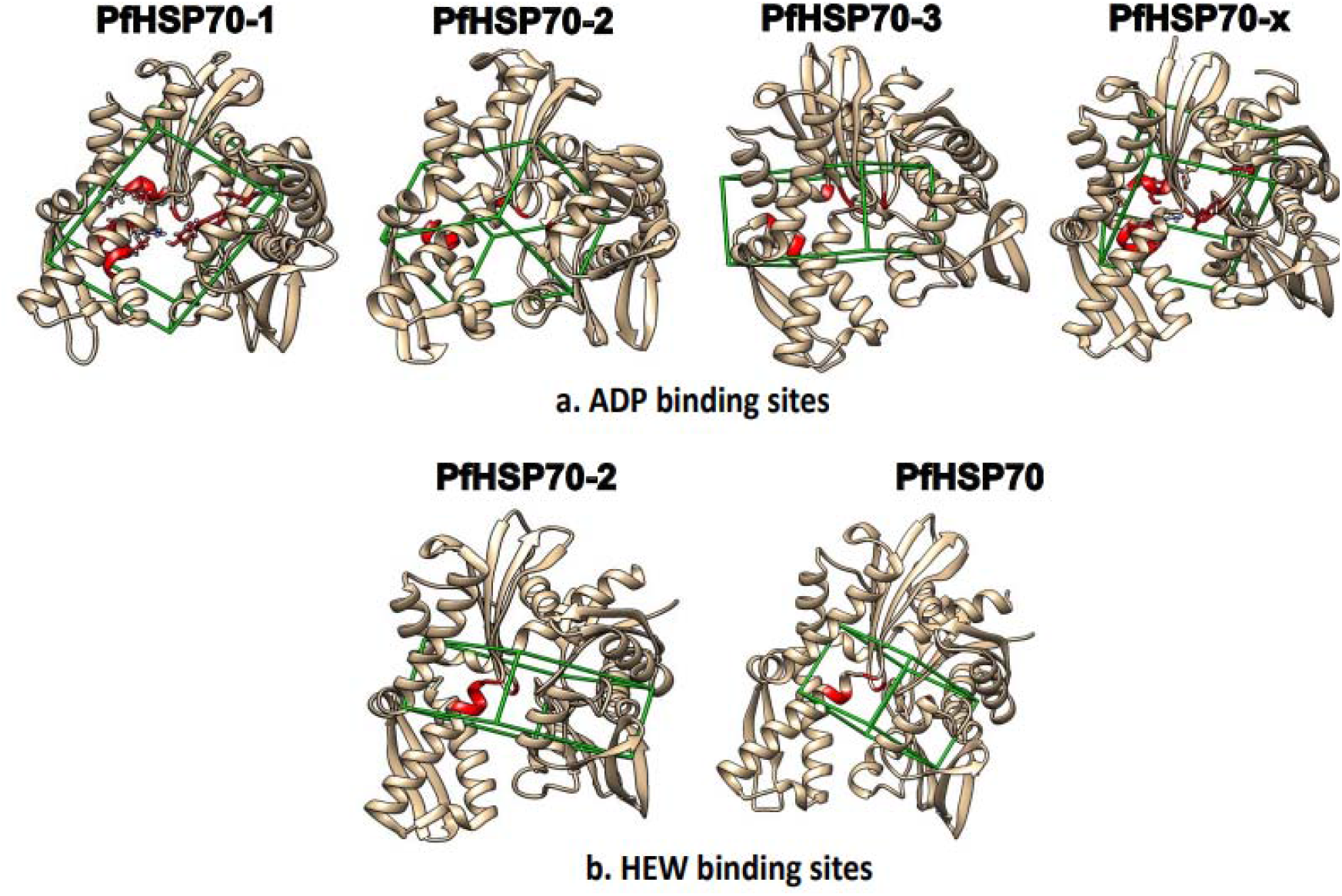
Grid boxes for in silico docking. Gridlines (green) are drawn encompassing ADP binding sites (red) on a. PfHSP70-1 b. PfHSP70-2 c. PfHSP70-3 d. PfHSP70-x and HEW binding sites (red) on e. PfHSP70-2 and f. PfHSP70-3. Protein molecules are shown as beige ribbon diagrams. Grid boxes for HEW binding sites on PfHSP70-1 and PfHSP70-x were same as Singh *et al* [13]

### Binding energies and prediction of YM-1 interacting residues

Binding energies computed from each docking experiment were used to select one binding pose each for PfHSP70-YM-1 complexes at either site (Fig. 3, Table 2). PfHSP70-3 displayed the highest affinity for YM-1 at the HEW binding site (−9.949). It is important to note that binding scores of the mitochondrial ‘PfHSP70-3’ for YM-1 at either binding site were significantly better than the other homologs (Table 2).

**Table 2:** Binding scores of PfHSP70-YM-1 docked complexes at ligand binding sites. Binding energies of respective PfHSP70s for YM-1 obtained from Chimera after docking are tabulated.

| Protein | Binding scores of ADP site | Binding scores of NCL site |
| --- | --- | --- |
| PfHSP70-1 | -6.155 | -6.38 |
| PfHSP70-2 | -7.05 | -4.747 |
| PfHSP70-3 | -9.165 | -9.949 |
| PfHSP70-x | -6.405 | -5.428 |

**Figure 3:**
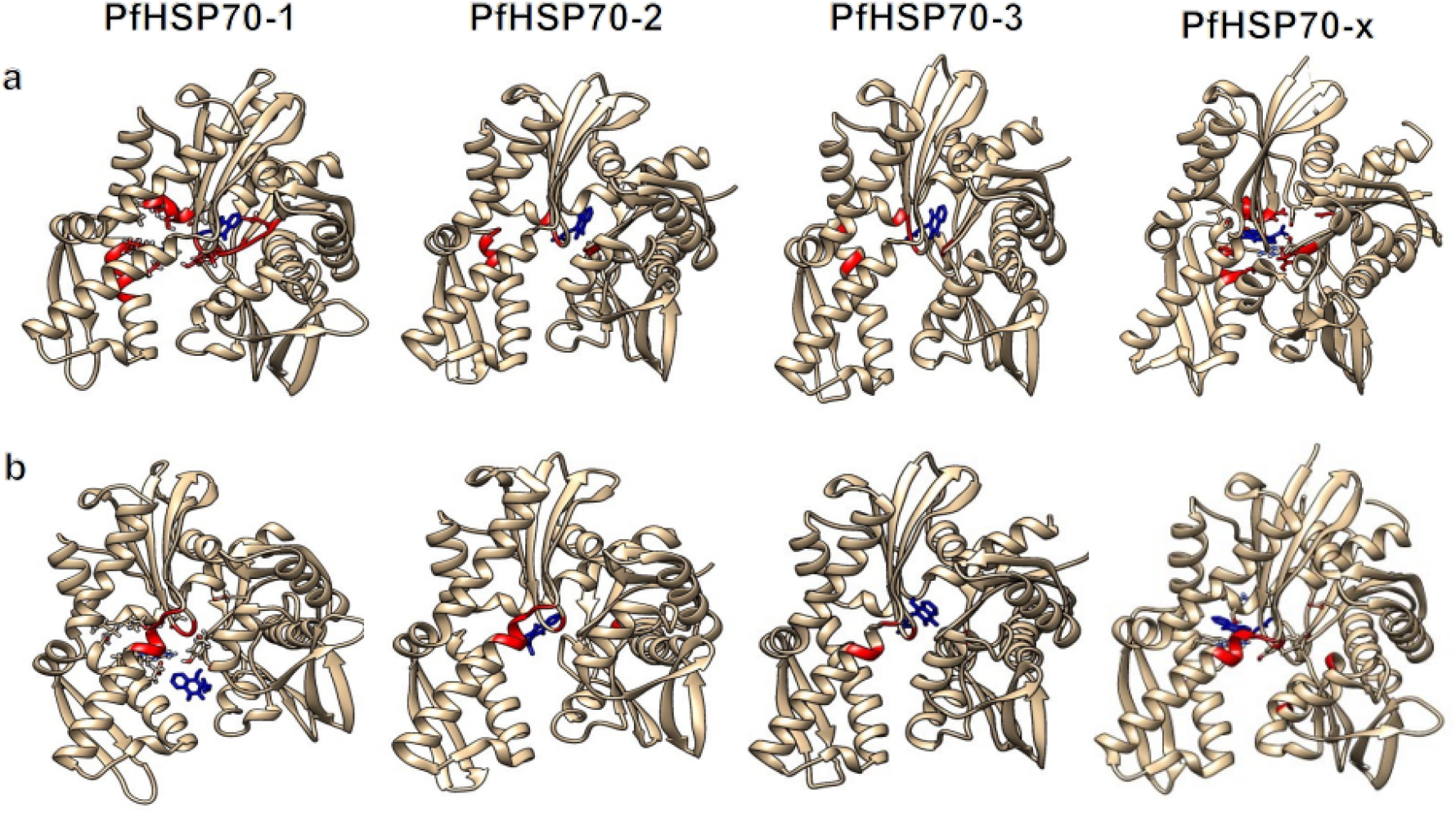
PfHSP70-YM-1 docked complexes. YM-1 docked on all PfHSP70 homologs (labelled). PfHSP70s are shown in beige. YM-1 (blue) docked at a-d: ADP binding site. e-h: HEW binding site.

The PfHSP70-3-YM-1 complex was analyzed by LigPlot to predict specific residues of the protein that are engaged in inhibitor binding. These are F66, L87, H223, E227 and D228. This interaction involves a total of five amino acids that bind YM-1 by making hydrophobic/ Van Der Waals contacts (Fig. 4). None of the residues were involved in hydrogen bonding.

**Figure 4:**
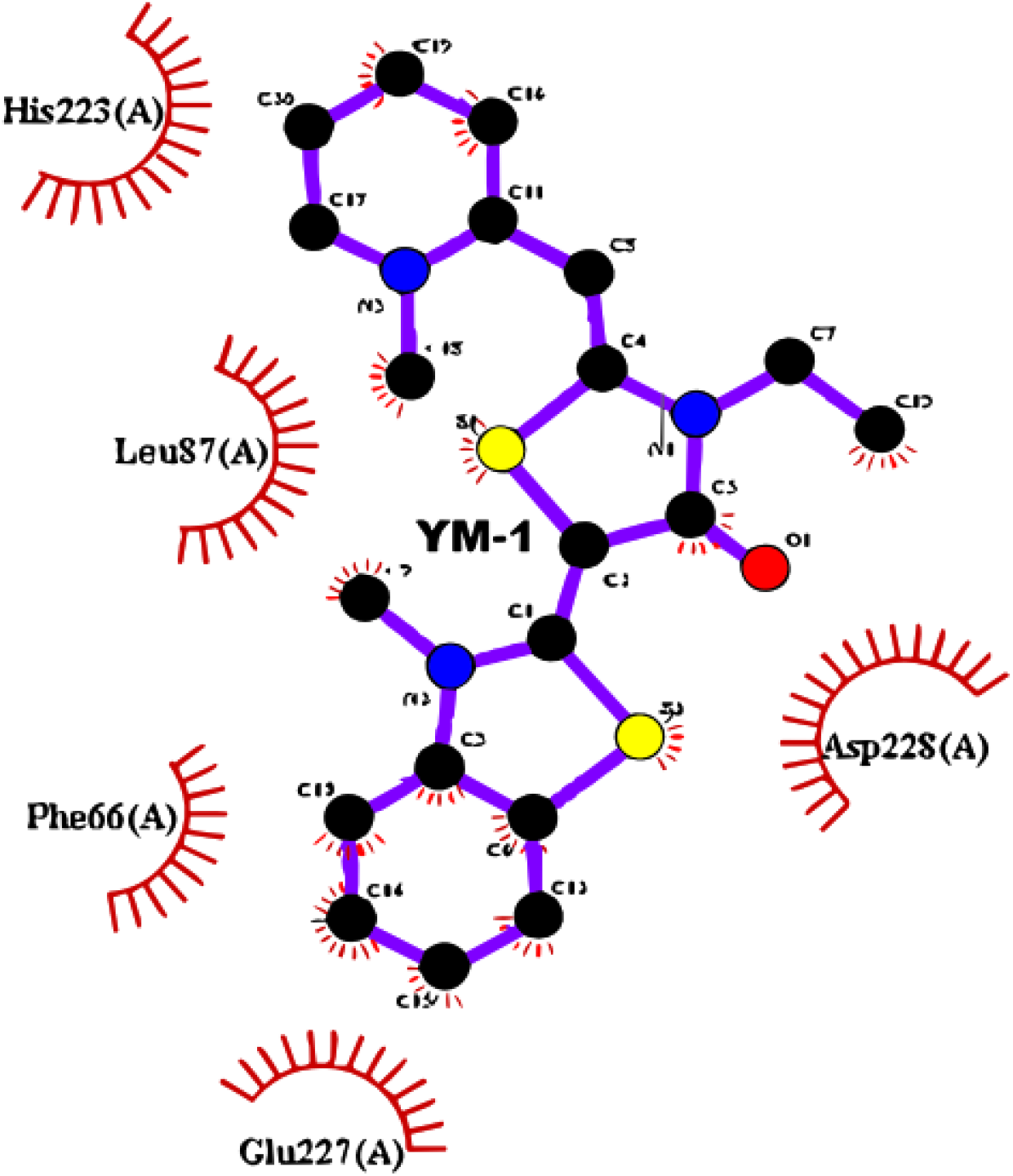
Interaction analysis using Ligplot for PfHsp70-3-YM-1 at HEW binding site. Ball and stick representation of YM-1 in the centre and interacting amino acids of PfHSP70-3 surrounding it are shown. All amino acids of the protein involved in non-bonding interactions (except hydrogen bonds) are depicted as spiked arcs.

## Conclusion

*Plasmodium falciparum*, the most common causative organism for fatal malaria expresses a family of HSP70s with four members with distinct subcellular localizations. YM-1 is a MKT-077 derivative that is being exploited for its activity against HSP70s to treat cancer. Since MKT-077 is effective against Pf cultures, YM-1 is likely to have similar ATPase inhibition activity by blocking binding of NBD with NEFs. Our bioinformatics approach has revealed that YM-1 engages preferentially with the HEW binding site of PfHSP70-1 (Binding energy: -6.38) and PfHSP70-3 (Binding energy: -9.949). On the other hand, PfHSP70-2 (Binding energy: -7.05) and PfHSP70-x (Binding energy: -6.405) have a better YM-1 binding score at the ADP site. HEW site of the mitochondrial PfHSP70-3 is predicted to have highest affinity for YM-1. Our analysis of interacting residues in the PfHSP70-3-YM-1 complex has helped us to predict amino acids that are involved in the ligand binding. These residues are involved in non-bonding interactions except hydrogen bonding. Information derived from the present study may form the foundation for design and development of newer rhodacyanine based antimalarials.

## Abbreviations

Pf: *Plasmodium falciparum*
HSP: Heat shock protein
SBD: Substrate binding domain
NBD: Nucleotide binding domain
ATP: Adenosine tri phosphate
ADP: Adenosine di phosphate
BAG3: Bcl-2-associated AnthanoGene-3
NEF: nucleotide exchange factor
HIP: Hsp70 interacting proteins

## Declaration

*The authors have no relevant financial or non-financial interests to disclose*.

The authors declare that there are no conflicts of interest.

*All authors read and approved the final manuscript*.

## Acknowledgments

CN and VU were receiving scholarship from DBT, Government of India. SK is a DBT-SRF. The laboratories of PCM and RH are funded by UGC-DAE, and were earlier funded by DBT, RUSA and DST, Government of India.

## Figure legends

**Table S1:** Gridlines for molecular docking. The size and centre positions for the selected gridlines around ADP and HEW binding pockets are tabulated for various PfHSP70 homologs. Grid box dimensions for HEW binding sites on PfHSP70-1 and PfHSP70-x are the same as Singh *et al* [13].

| Protein | Coordinates | Centre<br>(ADP) | Size<br>(ADP) | Cetre<br>(HEW) | Size<br>(HEW) |
| --- | --- | --- | --- | --- | --- |
| PfHSP70-1 | X | 28.324, | 20.0035, | 38.2619, | 21.0054, |
|  | Y | 79.1629, | 24.4346, | 76.706, | 24.7739, |
|  | Z | 58.552 | 31.9765 | 59.6672 | 20.2875 |
| PfHSP70-2 | X | 37.3722, | 24.6118, | 24.7038, | 26.7157, |
|  | Y | 2.61098, | 18.3328, | -3.76068, | 12.9878, |
|  | Z | 32.0131 | 20.5079 | 30.6289 | 25.9824 |
| PfHSP70-3 | X | 36.9638, | 27.947, | 24.914, | 20.4841, |
|  | Y | -3.1422, | 16.6676, | -3.71376, | 14.2106, |
|  | Z | 31.1086 | 19.5887 | 30.1192 | 18.9535 |
| PfHSP70-x | X | 24.2115, | 16.9036, | 33.0404, | 27.0616, |
|  | Y | 77.466, | 21.4862, | 76.92881, | 18.8902, |
|  | Z | 58.4413 | 28.1978 | 54.4074 | 14.6084 |

## References

[1] Z. Albakova, G. A. Armeev, L. M. Kanevskiy, E. I. Kovalenko, and A. M. Sapozhnikov, ‘HSP70 Multi-Functionality in Cancer’, Cells, vol. 9, no. 3, p. 587, Mar. 2020, doi: 10.3390/cells9030587.

[2] A. Rousaki, Y. Miyata, U. K. Jinwal, C. A. Dickey, J. E. Gestwicki, and E. R. P. Zuiderweg, ‘Allosteric drugs: the interaction of antitumor compound MKT-077 with human Hsp70 chaperones’, J. Mol. Biol., vol. 411, no. 3, pp. 614–632, Aug. 2011, doi: 10.1016/j.jmb.2011.06.003.

[3] K. Takasu et al., ‘Rhodacyanine dyes as antimalarials. 1. Preliminary evaluation of their activity and toxicity’, J. Med. Chem., vol. 45, no. 5, pp. 995–998, Feb. 2002, doi: 10.1021/jm0155704.

[4] R. Moradi-Marjaneh, M. Paseban, and M. Moradi Marjaneh, ‘Hsp70 inhibitors: Implications for the treatment of colorectal cancer’, IUBMB Life, vol. 71, no. 12, pp. 1834–1845, Dec. 2019, doi: 10.1002/iub.2157.

[5] S.-K. Hong, D. Starenki, O. T. Johnson, J. E. Gestwicki, and J.-I. Park, ‘Analogs of the Heat Shock Protein 70 Inhibitor MKT-077 Suppress Medullary Thyroid Carcinoma Cells’, Int. J. Mol. Sci., vol. 23, no. 3, p. 1063, Jan. 2022, doi: 10.3390/ijms23031063.

[6] J. Koren et al., ‘Rhodacyanine derivative selectively targets cancer cells and overcomes tamoxifen resistance’, PloS One, vol. 7, no. 4, p. e35566, 2012, doi: 10.1371/journal.pone.0035566.

[7] T. A. Colvin et al., ‘Hsp70–Bag3 Interactions Regulate Cancer-Related Signaling Networks’, Cancer Res., vol. 74, no. 17, pp. 4731–4740, Sep. 2014, doi: 10.1158/0008-5472.CAN-14-0747.

[8] E. R. P. Zuiderweg, L. E. Hightower, and J. E. Gestwicki, ‘The remarkable multivalency of the Hsp70 chaperones’, Cell Stress Chaperones, vol. 22, no. 2, pp. 173–189, Mar. 2017, doi: 10.1007/s12192-017-0776-y.

[9] M. Gamerdinger, P. Hajieva, A. M. Kaya, U. Wolfrum, F. U. Hartl, and C. Behl, ‘Protein quality control during aging involves recruitment of the macroautophagy pathway by BAG3’, EMBO J., vol. 28, no. 7, pp. 889–901, Apr. 2009, doi: 10.1038/emboj.2009.29.

[10] ‘Höfeld: Hip, a novel cochaperone involved in the… -Google Scholar’. https://scholar.google.com/scholar_lookup? (accessed Jul. 26, 2023).

[11] H. M. Berman et al., ‘The Protein Data Bank’, Acta Crystallogr. D Biol. Crystallogr., vol. 58, no. Pt 6 No 1, pp. 899–907, Jun. 2002, doi: 10.1107/s0907444902003451.

[12] J. Day, A. Passecker, H.-P. Beck, and I. Vakonakis, ‘The Plasmodium falciparum Hsp70-x chaperone assists the heat stress response of the malaria parasite’, FASEB J., vol. 33, no. 12, pp. 14611–14624, Dec. 2019, doi: 10.1096/fj.201901741R.

[13] B. Singh et al., ‘Understanding the structural basis for differential binding of lapachol with PfHSP70s’. bioRxiv, p. 2023.07.18.549604, Jul. 19, 2023. doi: 10.1101/2023.07.18.549604.

[14] PubChem, ‘PubChem’. https://pubchem.ncbi.nlm.nih.gov/ x(accessed Jul. 13, 2023).

[15] R. A. Laskowski and M. B. Swindells, ‘LigPlot+: multiple ligand-protein interaction diagrams for drug discovery’, J. Chem. Inf. Model., vol. 51, no. 10, pp. 2778–2786, Oct. 2011, doi: 10.1021/ci200227u.

[16] B. Sk, B. Hm, K. Gj, M. Jl, N. H, and V. S, ‘Protein Data Bank (PDB): The Single Global Macromolecular Structure Archive’, Methods Mol. Biol. Clifton NJ, vol. 1607, 2017, doi: 10.1007/978-1-4939-7000-1_26.

[17] T. A. Tatusova and T. L. Madden, ‘BLAST 2 Sequences, a new tool for comparing protein and nucleotide sequences’, FEMS Microbiol. Lett., vol. 174, no. 2, pp. 247–250, May 1999, doi: 10.1111/j.1574-6968.1999.tb13575.x.

[18] E. F. Pettersen et al., ‘UCSF Chimera--a visualization system for exploratory research and analysis’, J. Comput. Chem., vol. 25, no. 13, pp. 1605–1612, Oct. 2004, doi: 10.1002/jcc.20084.

[19] J. Eberhardt, D. Santos-Martins, A. F. Tillack, and S. Forli, ‘AutoDock Vina 1.2.0: New Docking Methods, Expanded Force Field, and Python Bindings’, J. Chem. Inf. Model., vol. 61, no. 8, pp. 3891–3898, Aug. 2021, doi: 10.1021/acs.jcim.1c00203.

[20] J. Amick et al., ‘Crystal structure of the nucleotide-binding domain of mortalin, the mitochondrial Hsp70 chaperone’, Protein Sci. Publ. Protein Soc., vol. 23, no. 6, pp. 833–842, Jun. 2014, doi: 10.1002/pro.2466.

[21] A. Waterhouse et al., ‘SWISS-MODEL: homology modelling of protein structures and complexes’, Nucleic Acids Res., vol. 46, no. W1, pp. W296–W303, 2018.

[22] Y. Chen, C. Murillo-Solano, M. G. Kirkpatrick, T. Antoshchenko, H.-W. Park, and J. C. Pizarro, ‘Repurposing drugs to target the malaria parasite unfolding protein response’, Sci. Rep., vol. 8, no. 1, p. 10333, Jul. 2018, doi: 10.1038/s41598-018-28608-2.

